# Dynamic enhancer partitioning instructs activation of a growth regulator during exit from naïve pluripotency

**DOI:** 10.1101/441824

**Authors:** Maxim V.C. Greenberg, Aurélie Teissandier, Marius Walter, Daan Noordermeer, Deborah Bourc’his

## Abstract

During early mammalian development, the genome undergoes profound transitions in chromatin states, topological organization and recruitment of *cis* regulatory factors involved in transcriptional control. How these three layers of gene regulation interact is the matter of intense research. The *Zdbf2* gene—which is involved in growth control—provides a valuable model to study this question: upon exit from naïve pluripotency and prior to tissue differentiation, it undergoes a switch in usage from a distal to a proximal promoter, along with a switch in chromatin states, from polycomb to DNA methylation occupancy. Using an embryonic stem cell (ESC) culture system to mimic this period, we show here that four enhancers contribute to the *Zdbf2* promoter switch, concomitantly with dynamic changes in chromosome architecture. Indeed, CTCF plays a key role in partitioning the locus in ESCs, to facilitate enhancer contact with the distal *Zdbf2* promoter only. Partition relieving enhances proximal *Zdbf2* promoter activity, as observed during differentiation or with mutants that lack local CTCF-based partition. Importantly, we show that CTCF-based regulation occurs independently of the polycomb and DNA methylation pathways. Our study reveals the importance of multi-layered regulatory frameworks to ensure proper spatio-temporal activation of developmentally important genes.

## INTRODUCTION

During the early stages of mammalian development, as the embryo implants into uterine wall, the pluripotent cells that will go on to form somatic tissues transition from “naïve” to “primed” for lineage specification (*1*). One hallmark of the naïve pluripotent state is globally low DNA methylation, whereas primed cells are highly DNA methylated (*2*). Incidentally, chromatin architecture and the underlying histone modification landscape are also dramatically remodeled during this period (*3, 4*). Collectively, this process is referred to as epigenetic reprogramming, and it accompanies dynamic changes to the transcriptional landscape.

How DNA methylation, chromatin regulators, and chromosome conformation all cross-talk during epigenetic reprogramming has not fully crystallized. For example, the polycomb-group (PcG) proteins are developmentally important regulators of transcriptional repression; DNA methylation and PcG proteins are typically mutually exclusive at CpG-rich regions of the genome, such as CpG Island (CGI) promoters (*5–8*). Moreover, polycomb complexes have been proposed to be nodes of long-distance genomic interactions (*9–11*). Therefore, it is conceivable that DNA methylation may impact chromatin architecture by influencing the polycomb landscape.

Mammalian genomes are physically subdivided into “regulatory neighborhoods” known as topologically associated domains (TADs), which average roughly one megabase in size (*12, 13*). The CCCTC-BINDING FACTOR (CTCF) is absolutely required for TAD formation, and is bound at the majority of TAD borders, restricting inter-TAD interactions (*14*). Incidentally, CTCF is DNA methylation sensitive at a large subset of binding sites (*15*). It has even been reported in certain cancers, that ectopic DNA methylation impacts CTCF-mediated insulation between topologically associated domains (TADs) (*16*).

However, recently studies have suggested that DNA methylation does not play a major role in TAD organization. Firstly, embryonic stem cells which harbor knockouts for three DNA methyltransferases (*Dnmt* tKO), and are completely devoid of DNA methylation (*17*), have a virtually identical TAD landscape as their wild-type (WT) counterparts (*18*). Secondly, the somatic TAD organization is mostly established prior to the blastocyst stage, before the *de novo* DNA methylation program (*19*). Therefore, if DNA methylation does play a role in chromatin interactions, it is only at the sub-megabase level. One classic example where this is case is at the imprinted *H19/Igf2* locus, where differential methylation of a CTCF-binding site impacts enhancer-promoter looping (*20*).

To assess if and how the differentiation program may influence dynamic chromosome architecture, we utilized the imprinted *Zdbf2* locus as a model, which provides an intriguing example of dynamic regulation during differentiation. In the naïve state on the paternal allele, the distal *Long isoform of Zdbf2* promoter (*pLiz*) is active. As cells exit naïve pluripotency, a promoter switch occurs, resulting in activation of the proximal *Zdbf2* promoter (*pZdbf2*), located 73 kilobases (kb) downstream. From the primed state and then throughout life, *pZdbf2* is the functional promoter while *pLiz* is constitutively silenced. It should be noted that despite the genomic distance, there is no change in the message between the *Liz* and *Zdbf2* isoforms, thus the promoter switch does not result in protein diversity. Rather, there is a stratified relationship between the two promoters: *pLiz* activity is absolutely required for deposition of DNA methylation at a somatic differentially methylated region (sDMR). The sDMR DNA methylation, in turn, antagonizes polycomb-mediated repression, freeing *pZdbf2* (Figure S1A) (*21, 22*). Mice that are deficient for *Liz* are never able to activate *Zdbf2*, and this leads to a substantial growth defect. Thus, the *Zdbf2* locus provides a valuable model to dissect how promoter switching occurs in concert with shifting chromatin dynamics during cellular differentiation.

We show here using a cell-based approach, that several enhancers cooperate to regulate the dynamics of *Liz* and/or *Zdbf2* promoter activity. Moreover, CTCF-CTCF contacts at the locus change dynamically during differentiation, and contributed to the proper activity of *pLiz* and *pZdbf2*. Saliently, CTCF appears to exert its effect epistatically with respect to the DNA methylation and PcG pathways. This implies that there are two largely independent layers of chromatin-based control of *Zdbf2:* one at the level of chromatin marks and the other at the level of chromatin architecture. The highly regulated nature of *Zdbf2* underscores the importance of structural chromosome topology occurring in concert with chromatin marks to control proper spatio-temporal expression of developmentally consequential genes.

## RESULTS

### Two Classes of Putative Enhancers Lie in the *Liz/Zdbf2* Locus

To discover functional genetic elements that regulate *Zdbf2* alternative promoter usage during the *de novo* DNA methylation program, we performed an assay for transposase-accessible chromatin followed by sequencing (ATAC-Seq) (*23*) in naïve ESCs (cultured in 2i/LIF+vitC), when *pLiz* is active and *pZdbf2* is repressed (*22*) (Figure 1A). We previously showed that by differentiating ESCs into primed, highly DNA methylated epiblast-like cells (EpiLCs), we can faithfully recapitulate *in vivo pLiz* to *pZdbf2* promoter usage dynamics (*22*). Therefore, we also performed ATAC-Seq in day 7 (D7) EpiLCs (Figure 1A). As expected, the ATAC-Seq peak for *pLiz* diminished as it became repressed and DNA methylated in EpiLCs. An ATAC-Seq peak was already present at *pZdbf2* in ESCs; this is correlated with our previous data indicating that *pZdbf2* is bivalent and poised in ESCs (*22, 24*).

**Figure 1.**
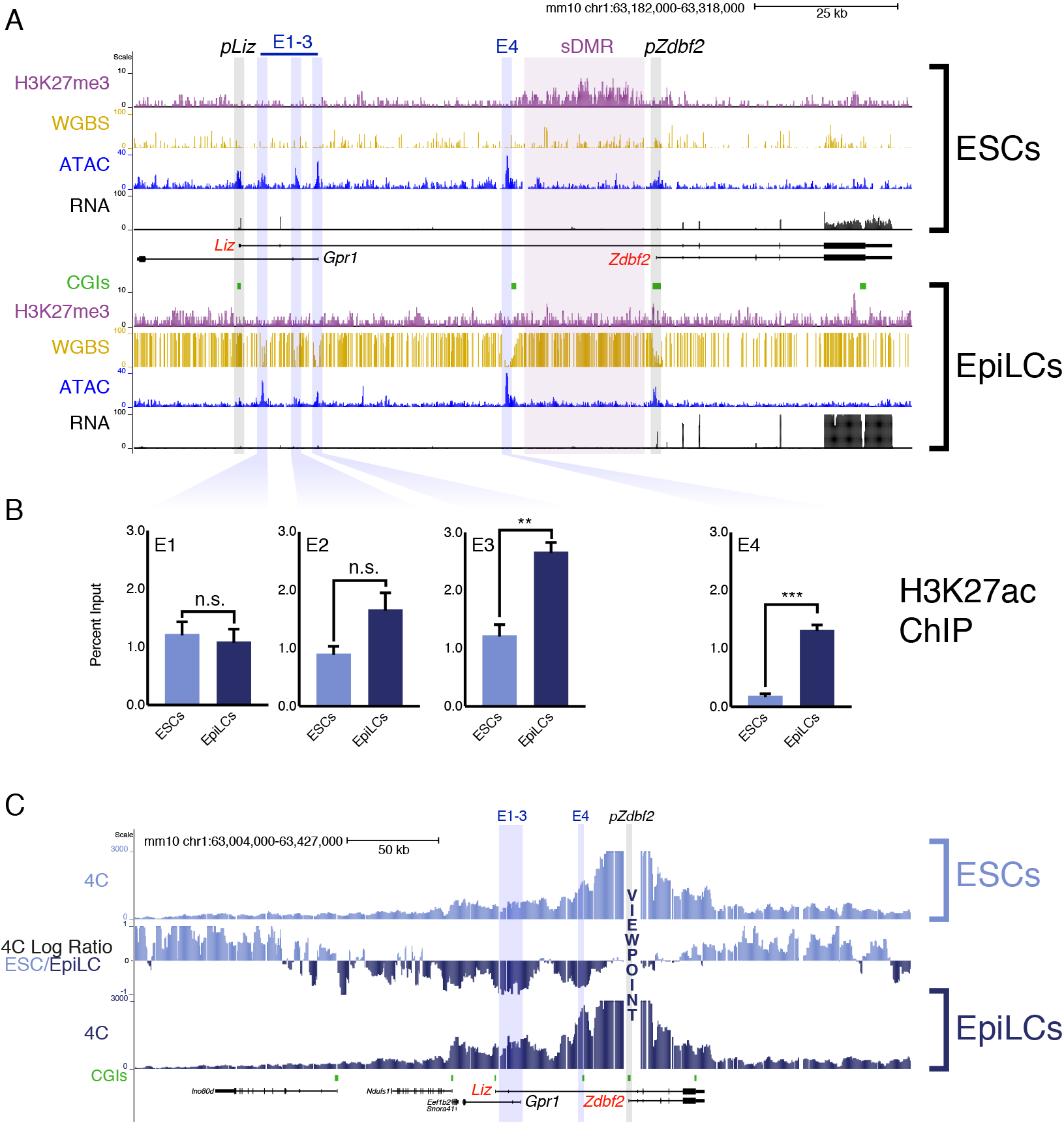
Chromatin dynamics of the *Liz/Zdbf2* locus during differentiation. **(A)** Chromatin and expression landscape in ESCs (top) and EpiLCs (bottom). In ESCs there is ~25kb block of H3K27me3 that extends through the *Zdbf2* promoter. The H3K27me3 signal depletes after DNA methylation is established. ATAC-seq reveals four prominent peaks of accessible chromatin between the *Liz* and *Zdbf2* promoters. The ATAC-seq peak at the *Liz* promoter decreases in EpiLCs concomitantly with decreased expression. WGBS: Whole genome bisulfite sequencing. H3K27me3 ChIP-seq data is from Greenberg et al., 2017. All other genomics data was generated for this study. One representative bioreplicate is displayed for each RNA-seq track. See also Figure S1. **(B)** H3K27ac ChIP-qPCR at the four inter-promoter ATAC-seq peaks. E1-3 are enriched for the mark in ESCs and EpiLCs, while E4 only becomes enriched in EpiLCs. Data is shown as ± s.e.m. from three biological replicates. See also Figure S1. **(C)** 4C-seq tracks from the *pZdbf2* VP in ESCs and EpiLCs. Ratios between 4C-seq signals is indicated in between the samples, and gene tracks are below. In EpiLCs, *pZdbf2* exhibits increased interactions at the four putative enhancers. The screen shot represents data from one biological replicate (two total). See also Figure S2. Statistical analyses were performed by two-tailed unpaired t-test: n.s = not significant, ** P ≤ 0.01, *** P ≤ 0.001.

In between the two promoters, four significant peaks were present in both ESCs and EpiLCs, three proximal to *pLiz* (E1-3), and one adjacent to a CGI that is an apparent border to the H3K27me3 block in ESCs (E4) (Figure 1A). Given that these regions of accessible chromatin were not lying on obvious active promoters, we reasoned that they were potential enhancer elements and named them E1 to E4, from the closest to the most distal to *pLiz*. Therefore we assayed for enrichment of H3K27 acetylation (H3K27ac), a mark of active enhancer elements (*25*). E1-3 appeared enriched for the H3K27ac mark in both cell types, while E4 was depleted for H3K27ac in ESCs and then became enriched for the mark in EpiLCs (Figure 1B). While E1-3 can be classified as active in ESCs and EpiLCs, the chromatin accessibility and H3K27ac dynamics at E4 are reminiscent of so-called “poised” enhancers (Figure S1B) (*26, 27*). Moreover, publicly available data indicates that E4 is marked by P300 in ESCs and shows high levels of vertebrate conservation (Figure S1C), two more features of poised enhancers (*26*).

The general regulator of pluripotency POU5F1/OCT4 is enriched at all of the putative enhancers in both naïve ESCs and EpiLCs (two pluripotent cell types), indicating that both classes of enhancers are likely regulated in a pluripotency-dependent manner (*27*). Importantly, publicly available *in vivo* data from the naïve pluripotent inner cell mass (ICM) of the blastocyst exhibit a chromatin accessibility and H3K27me3 pattern akin to what we observed in ESCs for the *Zdbf2* locus, suggesting that the *in vivo* and *in cellula* regulation are coherent (Figure S1C) (*28, 29*).

### High Resolution 4C Reveals Enhancer-Promoter Dynamic Interactions

Given that E1-4 exhibit the chromatin signature of enhancer elements, it is possible that the *Liz* and *Zdbf2* promoters undergo dynamic interactions with these regulatory elements during EpiLC differentiation. In order to test this, we performed high-resolution circular chromosome conformation capture followed by sequencing (4C-seq) during differentiation (*30, 31*). Available Hi-C data from mouse ESCs indicates that the *Zdbf2* locus exists within an “inter-TAD” that spans roughly 650kb (Figure S2A) (*32*). According to our 4C-seq data, this inter-TAD can be further subdivided, with intra-*Liz/Zdbf2* locus and intra-*Adam23* locus interactions occurring in relatively mutually exclusive domains (Figure S2B).

Using the *pLiz* as a 4C-seq viewpoint (VP), we did not observe distal looping that occurred at high frequency in either ESCs or EpiLCs (Figure S2C). However, in ESCs, when *Liz* is expressed, the promoter did exhibit increased interactions with the E1-3 cluster relative to EpiLCs, all of which are marked by H3K27ac in this cell type (Figure 1B). This is consistent with the possibility that E1-3 contribute to *Liz* regulation. Given the close proximity between *pLiz* and E1-3, it is not surprising that interactions remained high between them in EpiLCs, when *Liz* is repressed. No marked looping appeared to occur between *pLiz* and E4 in either ESCs or EpiLCs.

A clear picture emerged from the 4C-seq analysis for the *Zdbf2* promoter (*pZdbf2*) (Figure 1C). In EpiLCs, E1-4 were all marked by H3K27ac, and *pZdbf2* is active. Our 4C-seq revealed that in EpiLCs, *pZdbf2* exhibits increased contacts with all four of the putative enhancers, indicating a potential cooperative role for E1-4 in *Zdbf2* activation.

### Determination of Enhancer Function and Regulation

From our 4C-seq analyses we reasoned that E1-3 potentially regulate *pLiz* in ESCs, while E4 is silent (Figure 2A). To test this, we generated homozygous deletions of combinations of putative enhancer elements (Figure S2D). The E3 element also serves as the promoter of the *Gpr1* gene, which is lowly expressed in our system and we previously showed plays no role in *Zdbf2* regulation (*22*). As such, deleting the element had no impact on expression or DNA methylation at the *Zdbf2* locus (Figure S2E and S2F). If E3 is an enhancer element, it may be redundant with E1 and/or E2. Therefore, we generated a ~13kb deletion that encompassed E1-3 (Figure 2B and S2C). In the absence of these elements, the *Liz* transcript was markedly repressed, and the canonical *Zdbf2* isoform failed to properly activate (Figure 2C). As *Liz* is required to activate *Zdbf2*, it should be noted that this deletion does not confirm E1-3 elements regulate *pZdbf2*. However, the data provide a strong indication that E1-3 are indeed enhancers of *pLiz*.

**Figure 2.**
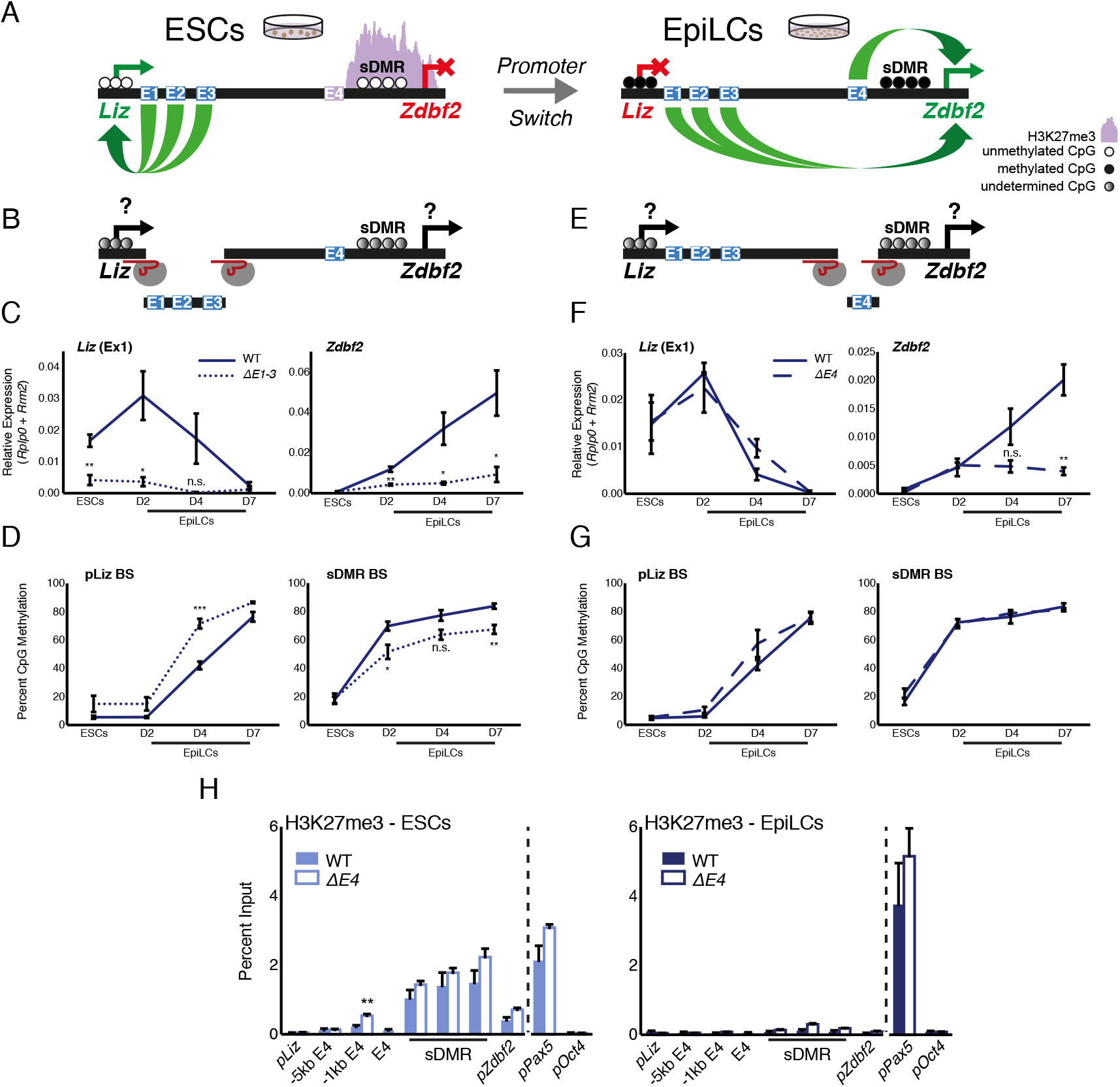
Genetic deletion analyses of enhancer elements. **(A)** Model for enhancer regulation based on 4C-seq data. See also Figure S2. **(B)** Model for deletion of E1-3. See also Figure S2. **(C)** RT-qPCR of *Liz* (left) and *Zdbf2* (right) during EpiLC differentiation in WT and the Δ*E1-3* mutant. The Δ*E1-3* mutation results in significant depletion in both transcripts. Data are shown as ± s.e.m. from five and three biological replicates for WT and mutant, respectively. **(D)** DNA methylation of *pLiz* (left) and the sDMR (right) during EpiLC differentiation as measured by bisulfite conversion followed by pyrosequencing (BS-pyro) in WT and the Δ*E1-3* mutant. When *Liz* fails to activate, DNA methylation is acquired faster at *pLiz*, and fails to properly accumulate at the sDMR. Data are shown as ± s.e.m. from five and three biological replicates for WT and mutant, respectively. **(E)** Model for deletion of E4. See also Figure S2. **(F)** RT-qPCR of *Liz* (left) and *Zdbf2* (right) during EpiLC differentiation in WT and the Δ*E4* mutant. There is no effect on *Liz* expression dynamics, but *Zdbf2* does not properly activate. Data are shown as ± s.e.m. from four biological replicates for each genotype. **(G)** DNA methylation of *pLiz* (left) and the sDMR (right) during EpiLC differentiation as measured by BS-pyro in WT and the Δ*E4* mutant. DNA methylation is unperturbed in the Δ*E4* mutant. Data are shown as ± s.e.m. from four biological replicates for each genotype. **(H)** H3K27me3 ChIP-qPCR in ESCs (left) and EpiLCs (right). There is no significant effect on polycomb dynamics in Δ*E4* mutation, except mild ectopic spreading upstream of the sDMR region. *pPax5* and *pOct4* are positive and negative controls, respectively. Data are shown as ± s.e.m. from three biological replicates for each genotype. Statistical analyses were performed by two-tailed unpaired t-test: n.s = not significant, * P ≤ 0.05, ** P ≤ 0.01, *** P ≤ 0.001.

We previously showed that DNA methylation accumulates at *pLiz* after transcription ablates (*22*). Interestingly, in the absence of E1-3, DNA methylation accumulated faster at *pLiz*, perhaps indicating less protection from *de novo* DNA methyltransferases due to reduced transcription factor occupancy (Figure 2D). *Liz* expression is required for DNA methylation establishment at the sDMR region *in cellula* and *in vivo* (*22*). It should be noted that in the cell-based system the imprint is lost and both alleles become methylated, but we maintain here the sDMR terminology. In the absence of E1-3, the DNA methylation failed to properly accumulate at the sDMR region, reaching 67% by D7 (Figure 2D). This was likely as a consequence of reduced *Liz* expression, as deletion of *pLiz* resulted in 45% sDMR methylation (*22*).

Upon the promoter switch, E4 became enriched for H3K27ac. We hypothesized that a deletion for E4 would have minimal impact on *pLiz*, but may affect *pZdbf2* activity (Figure 2A and 2E). Indeed, Δ*E4* mutant cells exhibited no alteration of *Liz* expression, but *Zdbf2* transcripts were strongly reduced (Figure 2F). As *Liz* was unaffected, there was no impact on DNA methylation at the locus (Figure 2G). Moreover, reduced expression of *Zdbf2* in Δ*E4* EpiLCs did not correlate with maintained polycomb occupancy in the sDMR region and *pZdbf2* (Figure 2H). In sum, the enhancer E4 is necessary for *pZdbf2* activation, regardless of the local DNA methylation or polycomb status.

The E4 enhancer element bears the hallmark of a poised enhancer in that it is enriched for P300 in ESCs, but only becomes active in EpiLCs. However, poised enhancers were originally defined as being enriched for H3K27me3 (*26*), whereas E4 is depleted for the mark (Figure 2H). H3K27me2, which like H3K27me3 is deposited by polycomb repressive complex 2 (PRC2), has also been reported to prevent firing of enhancers in ESCs (*33*). Yet H3K27me2 ChIP analysis revealed that E4 is depleted for this mark as well (Figure S3A). E4 does seem to play a role in preventing ectopic polycomb spreading: deleting E4 resulted in a slight increase of H3K27me3 enrichment 1kb upstream of the WT polycomb domain, however the signal was identical to WT levels by 5kb upstream (Figure 2H).

We previously showed that in ESCs containing loss-of-function mutations in the *Embryonic ectoderm development* (*Eed*) gene (*34*)—a core component of PRC2—there was precocious activation of *pZdbf2* (*22*). Therefore, we wanted to observe if a PRC2 mutant would result in a change in the chromatin status of E4. Indeed, both E4 and *pZdbf2* became enriched for H3K27ac in the absence of polycomb-mediated repression (Figure 3A). Incidentally, *pLiz* and E1-3, which are already active in ESCs, exhibited no significant change. In Δ*Liz* mutants, *Zdbf2* remains polycomb repressed (*22*). As such, in the Δ*Liz* mutant the E4 enhancer did not attain complete levels of H3K27ac during EpiLC differentiation (Figure 3B). In sum, while E4 does not display the signatures of direct polycomb regulation, *per se*, its activity is controlled in a polycomb-dependent manner.

**Figure 3.**
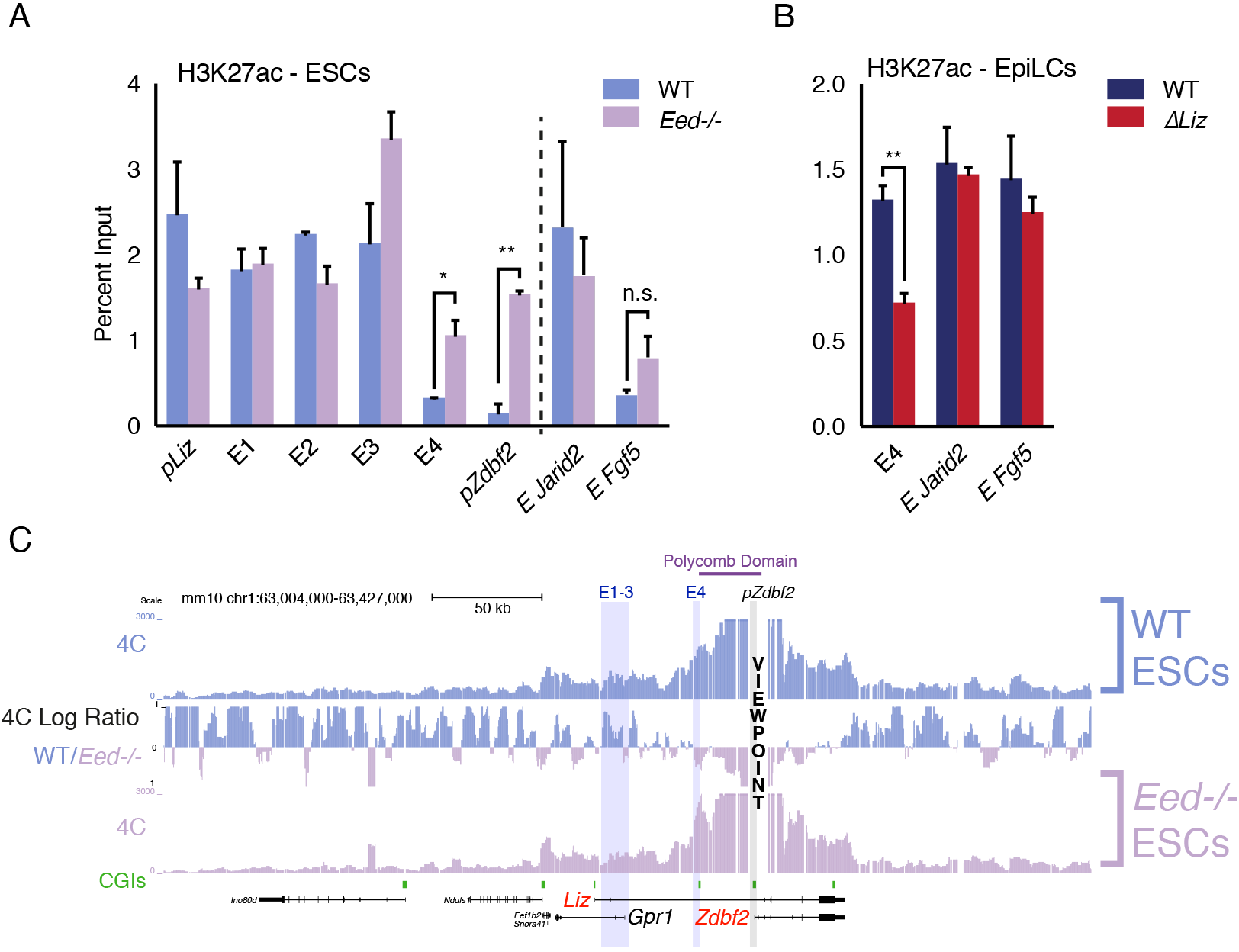
Polycomb regulates E4, but plays minor role in chromosome conformation. **(A)** H3K27ac ChlP-qPCR in WT and *Eed*−/− ESCs. *pLiz* and E1-3 are unaffected, but E4 and *pZdbf2* become aberrantly activated. Data are shown as ± s.e.m. from two biological replicates for both genotypes. **(B)** H3K27ac ChIP-qPCR in WT and Δ*Liz* EpiLCs. In Δ*Liz* mutants, when the sDMR remains enriched for H3K27me3, E4 remains diminished for H3K27ac. Data are shown as ± s.e.m. from three biological replicates for both genotypes. See also Figure S3. **(C)** 4C-seq tracks from the *pZdbf2* VP in WT and *Eed*−/− ESCs. Ratios between 4C-seq signals is indicated in between the samples, and gene tracks are below. In the polycomb mutant, *pZdbf2* exhibits increased interactions at E4, but not E1-3. See also Figure S3. Statistical analyses were performed by two-tailed unpaired t-test: n.s = not significant, * P ≤ 0.05, ** P ≤ 0.01.

### *Liz* Transcription and Polycomb Play a Minor Role in 3D Organization of the Locus

During differentiation, the transcription initiated from *pLiz* and traversing the locus is required for polycomb-to-DNA methylation switch, and *pZdbf2* activation (*22*). However, our 4C-seq analysis revealed that in the absence of *Liz* transcription, there is only a minor effect on the conformation of *pZdbf2* (Figure S3B). Moreover, in Δ*Liz* EpiLCs, *pZdbf2* exhibited increased interactions with E1-4, but not to the same extent as WT EpiLCs. It should be noted that the Δ*Liz* DNA methylation phenotype is only partial in the cell-based system, which may account for the intermediate chromosome conformation phenotype.

The polycomb region that regulates *pZdbf2* spans ~25kb, from E4 and into the body of *Zdbf2* (Figure 1A). Consistent with previous reports, in ESCs this region forms a tightly packed domain (*10*) (Figure 1C). We performed 4C-seq in *Eed* mutant ESCs in order to determine if polycomb impacts the chromosome conformation (Figure 3C). In fact, in a PRC2 mutant, *pZdbf2* interacted even more frequently within the polycomb domain. This is likely due to the activation of E4, and increased promoter-enhancer looping. It has recently been shown that active promoters exhibit increased agitation in the nucleus, leading to a potential increase of promoter-enhancer contacts (*35*). Given that *pZdbf2* becomes active in *Eed* mutant ESCs, logic would dictate that it would interact more frequently with E1-3, which are also active. However, our 4C-seq in the polycomb mutant showed that this was not the case (Figure 3C). To summarize, *Liz* transcription and the polycomb status play a limited role in the regulation of the *pZdbf2* interaction landscape, and there must be other mechanisms in place.

### CTCF Partitions the *Liz/Zdbf2* Locus in ESCs

Given that *pZdbf2* does not interact with E1-3 in polycomb mutant ESCs, those enhancers must be restricted from forming long-range loops. The most likely candidate to contribute to locus organization is CTCF (*36*). We analyzed the 4C-seq patterns of several CTCF binding sites (*37*) throughout the locus (data available upon request). In ESCs, a CTCF-binding site proximal to the *Gpr1* promoter formed a looping structure with two CTCF sites downstream of the *Gpr1* gene (Figure 4A). Incidentally, *pLiz* and E1-3 lie within this loop. During differentiation to EpiLCs, this looping structure was reduced. In accordance, the CTCF binding at this site was depleted, whereas CTCF remained bound at the sites downstream of *Gpr1* (Figure 4B). Therefore, this binding platform will be referred to as the “CTCF_partition site” (CTCF_PS), which physically separates the active *pLiz*/E1-3 region from the silent *pZdbf2/E4* in ESCs (Figure 4C). In EpiLCs, depletion of CTCF at the partition site would then allow for *pZdbf2* to interact with E1-3, while *pLiz* is silenced.

**Figure 4.**
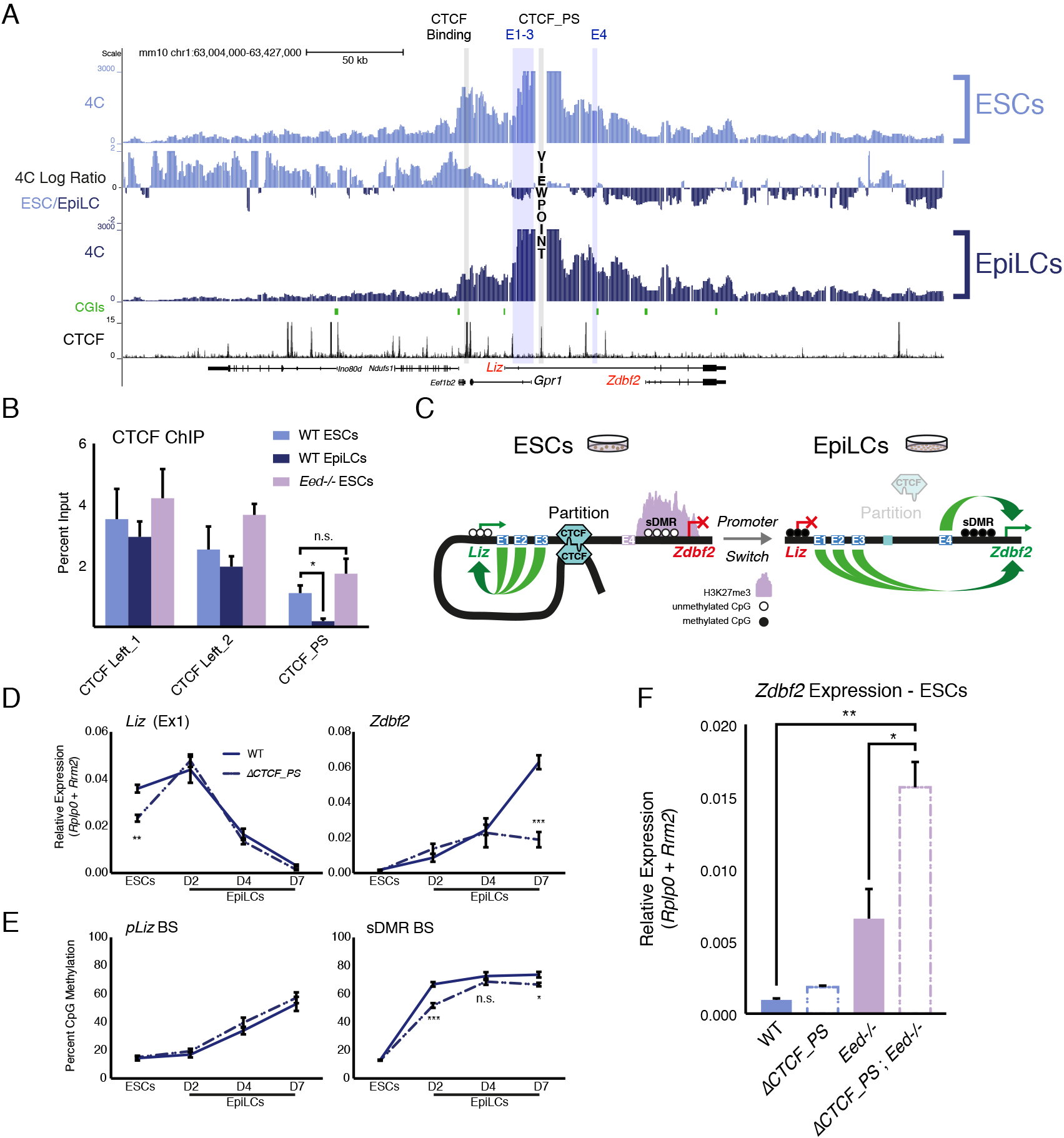
The *Liz/Zdbf2* locus is partitioned by CTCF, which instructs expression dynamics. **(A)** 4C-seq tracks from the CTCF_PS VP in WT ESCs and EpiLCs. Ratios between 4C-seq signals is indicated in between the samples, and gene and CTCF binding tracks (Stadler et al., 2011) are below. The CTCF_PS forms a loop with two CTCF sites downstream of the *Gpr1* gene. The looping diminishes in EpiLCs. See also Figure S3. **(B)** CTCF ChIP-qPCR in WT ESCs and EpiLCs and *Eed*−/− ESCs. CTCF binding remains unchanged on the two CTCF sites downstream of *Gpr1* (termed CTCF Left_1 and Left_2). At the CTCF_PS, CTCF binding is reduced in WT EpiLCs, which is correlated with decreased looping. CTCF binding remains enriched in *Eed*−/− ESCs, consistent with the maintained loop structure. Data are shown as ± s.e.m. from three biological replicates. See also Figure S4. **(C)** Model for CTCF-mediated partitioning of the locus. In ESCs, *pLiz* and E1-3 are active, and physically separated from the silent E4 and *pZdbf2*. During differentiation, the partition is diminished, allowing E1-3 to bolster *pZdbf2* activation, while *Liz* has become silent. **(D)** RT-qPCR of *Liz* (left) and *Zdbf2* (right) during EpiLC differentiation in WT and the Δ*CTCF_PS* mutant. *Liz* is less expressed in mutant ESCs, but reaches WT levels of expression during differentiation. Nevertheless, *Zdbf2* does not properly activate in the mutant. Data are shown as ± s.e.m. from four biological replicates for each genotype. See also Figure S4. **(E)** DNA methylation of *pLiz* (left) and the sDMR (right) during EpiLC differentiation as measured by BS-pyro in WT and the Δ *CTCF_PS* mutant. DNA methylation is unperturbed in the mutant at *pLiz*, but is reduced at the sDMR. Data are shown as ± s.e.m. from four biological replicates for each genotype. See also Figure S4. **(F)** RT-qPCR of *Zdbf2* in ESCs in absence of CTCF partition and/or PRC2. While *Zdbf2* is already upregulated in the *Eed* mutant, this effect is exacerbated in the absence the partition, likely because *pZdbf2* is less restrained from interacting with E1-3. Data are shown as ± s.e.m. from three biological replicates for each genotype. Statistical analyses were performed by two-tailed unpaired t-test: n.s. = not significant, * P ≤ 0.05, ** P ≤ 0.01, *** P ≤ 0.001.

Using the CTCF_PS as a VP in our *Eed* mutant ESCs, we observed that the partition loop still formed (Figure S3C). Furthermore, CTCF still remained enriched at the CTCF_PS in PRC2 mutant ESCs (Figure 4B). The continued formation of the partition in the absence of polycomb-mediated regulation would explain why *pZdbf2* failed to exhibit increased interactions with E1-3, even though the promoter has adopted an active state.

Given that CTCF is DNA methylation sensitive at a subset of binding sites (*15*), we reasoned that perhaps *de novo* DNA methylation is required for evicting CTCF from the CTCF_PS. We tested this by differentiating *Dnmt* tKO ESCs, which are able to differentiate to a state akin to WT EpiLCs despite a total lack of DNA methylation (*22, 38*). However, even in the absence of DNA methylation, CTCF depleted at the partition site (Figure S3D). A recent study reported that transcription can disrupt CTCF binding and chromatin architecture (*39*), yet we observed reduced CTCF enrichment even in the absence of the *Liz* transcript (Figure S3D). Therefore, the CTCF depletion at the CTCF_PS in EpiLCs is differentiation dependent, but independent of DNA methylation- or *Liz* transcription-based regulation.

### CTCF Partitioning Fine-tunes *pLiz* Programming of *pZdbf2*

To assess the regulatory impact of CTCF partitioning, we generated a deletion of the CTCF_PS (Figure S4A). 4C-seq in Δ*CTCF_PS* ESCs revealed that *pLiz* interacts less frequently with E1-3 (Figure S4B), perhaps as the promoter is less constrained without the CTCF partition. As such, *Liz* failed to properly express (Figure 4D). During differentiation, the *Liz* transcript was still able to attain WT levels, nevertheless *Zdbf2* failed to properly activate (Figure 4D). Moreover, while DNA methylation occurred normally at *pLiz*, the sDMR remained relatively hypomethylated compared to WT (Figure 4E), likely due to the delayed kinetics of *Liz* upregulation. Furthermore, the relative reduction of DNA methylation at the sDMR in Δ*CTCF_PS* mutant EpiLCs was correlated with a slight retention of H3K27me3 in comparison with WT, which may contribute to the failure of*pZdbf2* to properly activate (Figure 4D and S4C).

In Δ*CTCF_PS* mutant ESCs, *pZdbf2* exhibited increased contacts with E2 and E3 (Figure S4B). However, *Zdbf2* remained repressed, as polycomb enrichment remained unperturbed (Figure S4C). Given that in PRC2 mutant ESCs, *Zdbf2* is already partially de-repressed, we reasoned that by generating an *Eed* mutation in combination with deleting the partition site, we could observe further increase in *Zdbf2* expression, as *pZdbf2* would be unhindered from interacting with all four enhancers (Figure S4D). In parallel, we generated a new *Eed* mutation, so all cell lines would be in the identical genetic background (Figure S4D). We confirmed that the *Eed* mutant lines failed to exhibit EED protein nor H3K27me3 by western blotting (Figure S4E). Moreover, they showed no signs of precocious differentiation to EpiLCs (Figure S4F). Indeed, while *Zdbf2* was upregulated in absence of *Eed* alone, the expression was significantly increased in the Δ*CTCF-PS; Eed*-/- double mutant (Figure 4F). Therefore, we concluded that in addition to contributing to proper *pLiz* activation, the CTCF partition acts as a second level of protection, along with polycomb, to restrain precocious *pZdbf2* firing.

## DISCUSSION

In this study, we revealed the dynamic chromosome conformation of the *Liz*/*Zdbf2* locus that occurs concomitantly with epigenetic programming during differentiation. For proper *Zdbf2* activation, it is imperative to properly control *Liz* expression at the time *de novo* DNA methylation occurs. Here we show that a CTCF-structured loop organization forms in naïve ESCs. This partition allows for proper regulation of *pLiz,* which in turn can facilitate the epigenetic switch through *Liz* transcription. During differentiation the partitioning is relaxed, as *pLiz* becomes shut down, thus allowing distal enhancers to bolster *pZdbf2* activation. Deleting the CTCF_PS resulted in reduced *Liz* activation kinetics, but a fairly substantial effect on *Zdbf2* expression. This is in line with our previously published *pLiz* transcriptional interruption line, where *Liz* expression and DNA methylation are only moderately affected, but nonetheless *Zdbf2* cannot attain WT levels of activation (*22*). Such results underscore the sensitivity of *pZdbf2* activity to proper epigenetic programming.

In Δ*CTCF_PS* ESCs, *Zdbf2* transcription did not ectopically occur, even though there were no longer apparent restrictions for *pZdbf2* to interact with the active enhancers E1-3. The explanation for this is the polycomb-mediated silencing that persists over *pZdbf2* in the absence of CTCF_PS. In fact, the data suggest that the dynamics of the CTCF/partition axis and the polycomb/DNA methylation axis are decoupled at the *Zdbf2* locus. Thus, there are at least two layers of regulation of *Zdbf2* activation that act independently from classical transcription factor control at gene promoters: 1) instructive chromosome conformation allowing for proper *pLiz* and *pZdfb2* activity, and 2) a Liz-dependent epigenetic switch to evict polycomb at *pZdbf2.* This hierarchical model emphasizes the exquisite choreography that can occur to program developmentally important genes during the exit from the naïve pluripotent state.

One outstanding question is what is the cue that releases CTCF from the partition site during differentiation? The obvious candidate was DNA methylation, and while CTCF binding at the site may indeed be DNA methylation-sensitive, we found that CTCF was depleted during differentiation, whether the methyl mark was present or not. Another likely explanation is that CTCF is bound in combination with pluripotency-associated transcription factors. While CTCF is generally reported to be largely invariant across mammalian cell types (*40*), there are cell type-specific CTCF binding patterns, and roughly 60% occur in a DNA methylation-independent manner (*15*). For example, TATA BINDING PROTEIN ASSOCIATED FACTOR 3 (TAF3) is highly expressed in mouse ESCs, and mediates chromosome looping in concert with CTCF to regulate germ layer specification upon differentiation (*41*). While tAf3 does deplete in certain ESC differentiation protocols, at the RNA level it remains highly expressed in EpiLCs (RNA-seq from this study), which suggests it is not the factor controlling the partition at the *Liz/Zdbf2* locus. However, future experiments will be needed to test this possibility.

Our study presents a rare description of two alternative promoters utilizing a shared set of enhancers, but in a CTCF-guided, developmentally timed manner. In ESCs, CTCF is required to facilitate one promoter’s interactions (*pLiz*) while restricting the other’s (*pZdbf2*). Only upon removal of CTCF binding, the opportunity for *pZdbf2* is created to contact its enhancers. On its face, such a mechanism resembles the imprinted *Igf2/H19* locus, where CTCF binding near the *H19* gene on the maternal allele restrict interactions between a shared set of enhancers and the more distal *Igf2* gene (*42*). The *Igf2/H19* locus behaves differently than the *Liz/Zdbf2* locus, though, as differential CTCF binding is DNA methylation-dependent and set in the gametes—not dynamically regulated during cellular differentiation. Moreover, *Igf2* and *H19* are two different genes, not isoforms of the same gene.

CTCF has been shown to mediate enhancer switching. For example, at the *Hoxd* locus in mice, CTCF facilitates the interactions between the same set of gene promoters but different sets of enhancers depending on the context (*43*). Pertinently, dynamic enhancer switching during the naïve-to-primed differentiation is a common mechanism in mammals to maintain gene expression during this cellular transition (*44*); notably, the key pluripotency regulator *Oct4* relies on an ESC- and EpiLC-specific enhancers (*45, 46*). However, such an example represents an inverse scenario from *Zdbf2*, as the *Oct4* promoter remains unchanged.

Promoter switching is a widespread and developmentally important phenomenon (*47*). Aberrant promoter usage is associated with pathologies, such as cancer (*48*). CTCF is a likely candidate to organize genic three-dimensional structure and protect from aberrant promoter firing, as it does at *Zdbf2*. The *Zdbf2* locus presents a compelling case, because if *pLiz-to-pZdbf2* promoter switch does not occur at the proper developmental time, synchronized with the *de novo* DNA methylation program, *pZdbf2* cannot be activated (*22*). Notably, the organization of this locus is conserved between mouse and humans, implying the likelihood of a shared regulatory mechanism (*21*). Future studies should continue to shed light on the role that CTCF dynamics play in programming developmentally important promoter activity in the crucial window that precedes somatic tissue formation.

## ACKNOWLEDGMENTS

We would like to thank members of the Bourc’his laboratory and Nicolas Servant (Institut Curie) for insightful experimental and conceptual input. We thank Vincent Piras for sharing reanalyzed Hi-C matrices. We acknowledge the high-throughput sequencing facility of I2BC for its sequencing and bioinformatics expertise and for its contribution to this study for the 4C-seq (https://www.i2bc.paris-saclay.fr/spip.php?article399). The ICGex NGS platform of the Institut Curie (WGBS, ATAC-Seq) was supported by grants from the ANR-10-EQPX-03 (Equipex) and ANR-10-INBS-09-08 (France Génomique Consortium) from the Agence Nationale de la Recherche (″Investissements d’Avenir″ program), by the Canceropole Ile-de-France and by the SiRIC-Curie program-Grant ″INCa-DGOS-4654. The laboratory of D.B. is part of the Laboratoire d’Excellence (LABEX) entitled DEEP (11-LBX0044). This research was supported by the ERC (grant ERC-Cog EpiRepro). M.V.C.G. was supported by ARC and EMBO (LTF 457-2013) postdoctoral fellowships.

## AUTHOR CONTRIBUTIONS

Conceptualization, M.V.C.G., D.N., and D.B.; Methodology, M.V.C.G., D.N. and D.B.; Formal Analysis, A.T.; Investigation, M.V.C.G. and M.W.; Writing – Original Draft, M.V.C.G.; Writing – Review & Editing, M.V.C.G., D.N., and D.B.; Funding Acquisition, D.B.

## STAR ⋆ METHODS

### KEY RESOURCES TABLE

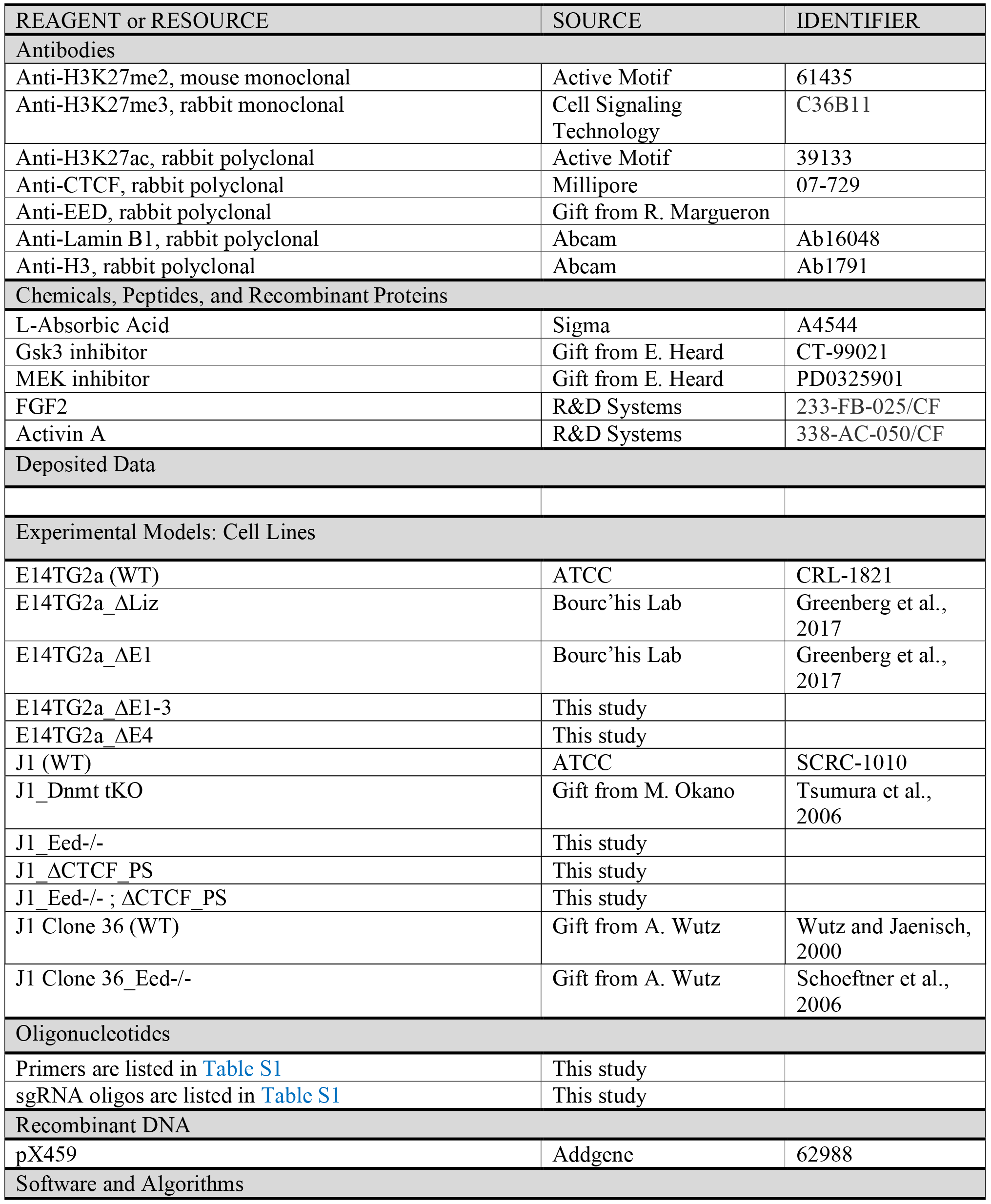

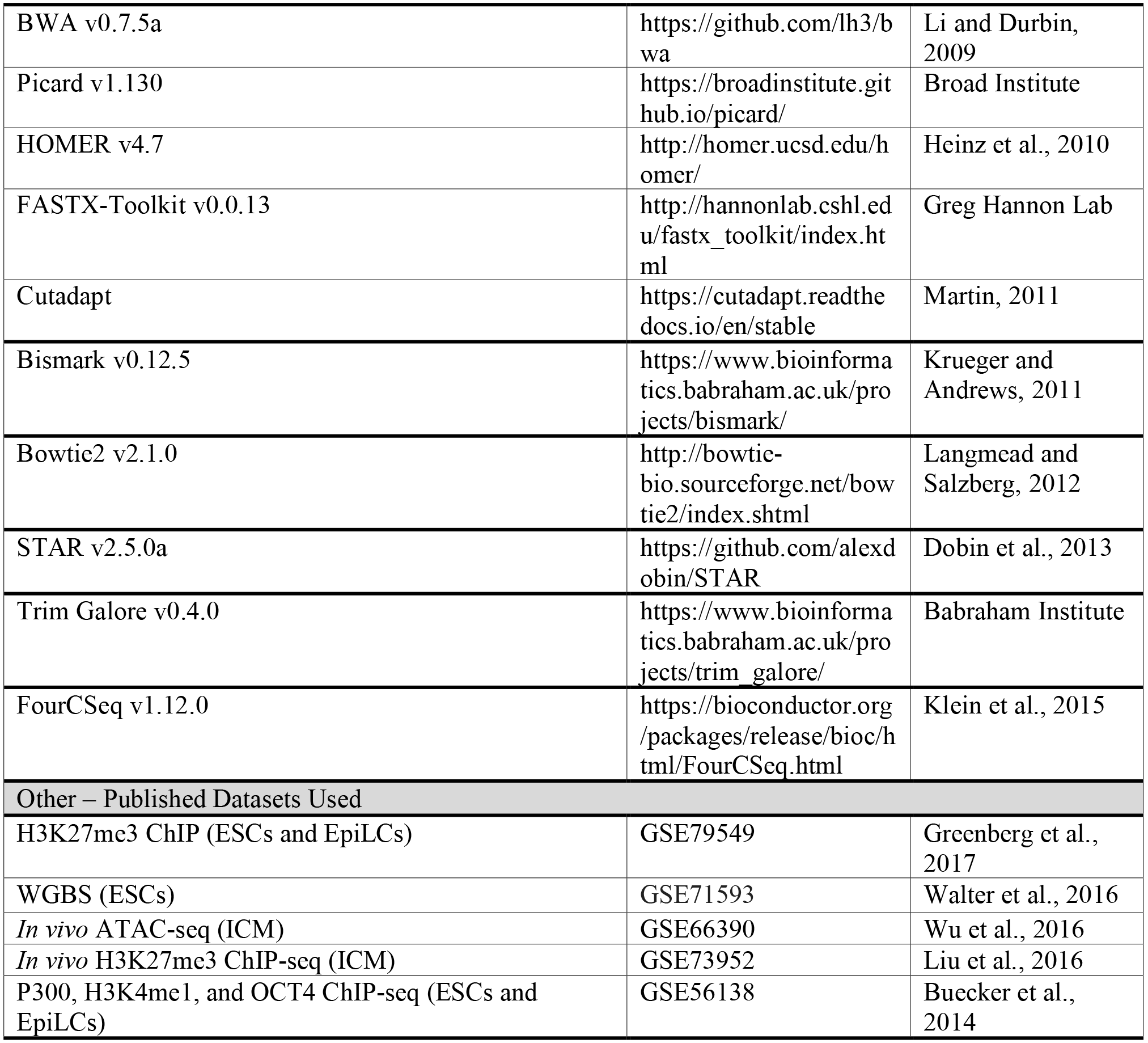

## EXPERIMENTAL MODEL AND SUBJECT DETAILS

### ESC Lines

All cell lines are listed in the Key Resource Table. For all experiments, the parental WT line was used as a control for mutant lines generated in that background. 4C-Seq was performed using clone 36 *Eed*−/− cells (J1 background). Therefore, when generating an in-house *Eed*−/− line, we used the same genetic background for consistency.

## METHOD DETAILS

### Cell culture and differentiation

Feeder-free ESCs were grown on gelatin-coated flasks. Serum culture conditions were as follows: Glasgow medium (Sigma) supplemented with 2mM L-Glutamine (Gibco), 0.1mM MEM non-essential amino acids (Gibco), 1mM sodium pyruvate (Gibco), 15% FBS, 0.1mM β-mercaptoethanol and 1000 U/ml leukemia inhibitory factor (LIF, Chemicon). Cells were passaged with trypsin replacement enzyme (Gibco) every two days. 2i culture conditions were as follows: N2B27 medium (50% neurobasal medium (Gibco), 50% DMEM/F12 (Gibco), 2mM L-glutamine (Gibco), 0.1mM β- mercaptoethanol, NDiff Neuro2 supplement (Millipore), B27 serum-free supplement (Gibco)) supplemented with 1000U/ml LIF and 2i (3 μM Gsk3 inhibitor CT-99021, 1 μM MEK inhibitor PD0325901). Cells were passaged every 2-4 days with Accutase (Gibco). Vitamin C (Sigma) was added at a final concentration of 100 μg/ml.

To induce EpiLC differentiation, cells were gently washed with PBS, dissociated, and replated at a density of 2^5^ cells/cm^2^ on Fibronectin (10 μg/ml)-coated plates in in N2B27 medium supplemented with 12 ng/ml Fgf2 (R&D) and 20ng/ml Activin A (R&D). EpiLCs were passaged with Accutase at D4 of differentiation.

Cells were regularly tested for presence of mycoplasma by sending used media to GATC/Eurofins for analysis.

### Generation of edited ESCs

All deletions in this study were generated with two CRISPR single guide RNAs (sgRNAs) specific to the target sequences followed by Cas9 nuclease activity and screedning for non-homologous end joining. sgRNAs were designed using the online CRISPOR online program (crispor.tefor.net) and cloned into the pX459 plasmid harboring the *Cas9* gene. All sgRNA sequences are listed in Table S1. Around five million WT serum-grown ESCs were transfected with 1-3μg of plasmid(s) using Amaxa 4d Nucleofector (Lonza) and plated at a low density. Ninety-six individual clones were picked and screened by PCR. Mutated alleles were confirmed by Sanger sequencing of cloned PCR amplicons. In the case of the *Eed* mutation, loss-of-function was further confirmed by immunoblotting.

### DNA methylation analyses

Genomic DNA from cells was isolated using the GenElute Mammalian Genomic DNA Miniprep Kit (Sigma), with RNase treatment.

Bisulfite conversion was performed on 500-1000ng of DNA using the EpiTect^®^ Bisulfite Kit (Qiagen). Bisulfite-treated DNA was PCR amplified and analyzed by pyrosequencing. Pyrosequencing was performed on the PyroMark Q24 (Qiagen) according to the manufacturer’s instructions, and results were analyzed with the associated software. All bisulfite primers are listed in Table S1. Statistical analyses were performed by a two-tailed unpaired *t-test* using GraphPad Prism6 software.

WGBS data from ESCs were previously generated (*10*) and EpiLCs were prepared from 50ng of bisulfite-converted genomic DNA using the EpiGnome/Truseq DNA Methylation Kit (Illumina) following the manufacturer instructions. Sequencing was performed in 100pb paired-end reads at a 30X coverage using the Illumina HiSeq2000 platform.

### ATAC-seq

ATAC-Seq was performed as described in Buenrostro et al., 2015 with minor modifications. Briefly, 50,000 cells were washed, but not lysed. Cells were transposed using the Nextera DNA library prep kit (Illumina) for 30 min at 37°. DNA was immediately purified using Qiagen MinElute Kit (Qiagen). qPCR was used to determine the optimal cycle number for library amplification. The libraries were sequenced on the Illumina HiSeq2500 platform to obtain 2×100 paired-end reads.

### Chromatin immunoprecipitation (ChIP)

ChIP was performed exactly as described in Walter et al., 2016. Briefly, cells were cross-linked directly in 15cm culture plates with 1% formaldehyde. After quenching with 0.125 M glycine, cells were washed in PBS and pelleted. After a three-step lysis, chromatin was sonicated with a Bioruptor (Diagenode) to reach a fragment size averaging 200 bp. Chromatin corresponding to 10 μg of DNA was incubated rotating overnight at 4°C with 3–5 μg of antibody. A fraction of chromatin extracts (5%) were taken aside for inputs. Antibody-bound chromatin was recovered using Protein G Agarose Columns (Active Motif). The antibody-chromatin mix was incubated in the column for 4 hr, washed eight times with modified RIPA buffer.

Chromatin was eluted with pre-warmed TE-SDS (50mM Tris pH 8.0, 10 mM EDTA, 1% SDS). ChlP-enriched sample and inputs were then reverse cross-linked at 65°C overnight and treated with RNase A and proteinase K. DNA was extracted with phenol/chloroform/isoamyl alcohol, precipitated with glycogen in sodium acetate and ethanol and finally resuspended in tris-buffered water. Enrichment compared to input was analyzed by qPCR using the Viia7 thermal cycling system (Applied Biosystems). Primers are listed in Table S1.

### 4C-seq

The design of VPs and preparation of 4C-seq libraries was performed as described in detail by Matelot and Noordermeer, 2016, with only minor modifications. DpnII or its isoschiszomer MboII (New England Biolabs) were chosen as the primary restriction enzyme, and NlaIII (New England Biolabs) as the secondary restriction enzyme. ESC and EpiLC material were harvested from 150cm^2^ culture flasks (TPP Techno Plastic Products AG), which provided ample material for up to four technical replicates presuming cells were healthy and near confluency. To avoid technical artifacts, crosslinking and library preparation were performed in parallel for each experiment. For each VP, approximately 1μg of library material was amplified using 16 individual PCR reactions with inverse primers containing indexed Illumina TruSeq adapters (primer sequences are listed in Table S1). PCR products were originally purified using the MinElute PCR purification kit (Qiagen) to remove unincorporated primer, but we found that purification was more efficiently performed using Agencourt AMPure XP beads (Beckman Coulter). Sequencing was performed on the Illumina NextSeq 500 system, using 75bp single-end reads with up to 14 VPs multiplexed per run.

### RNA expression

Total RNA was extracted using Trizol (Life Technologies), then DNase-treated and column purified (Qiagen RNeasy Kit). To generate cDNA, purified RNA was reverse transcribed with SuperscriptIII (Life Technologies) primed with random hexamers. RT-qPCR was performed using the SYBR Green Master Mix on the Viia7 thermal cycling system (Applied Biosystems). Relative expression levels were normalized to the geometric mean of the Ct for housekeeping genes *Rrm2* and *Rplp0* with the ΔΔCt method. Primers are listed in Table S1. Statistical analyses were performed by a two-tailed unpaired t-test using GraphPad Prism6 software.

RNA-seq libraries were prepared from 500ng of DNase-treated RNA with the TruSeq Stranded mRNA kit (Illumina). Sequencing was performed in 100pb paired-end reads using the Illumina HiSeq2500 platform.

### Immunoblotting

Western blots were visualized using the ChemiDoc MP (Biorad). The antibodies are listed in the Key Resource Table.

## QUANTIFICATION AND STATISTICAL ANALYSIS

### ATAC-seq analysis

2×100bp paired-end reads were aligned onto the Mouse reference genome (mm10) using Bwa mem v0.7.5a (*2*)with default parameters. Duplicate reads were removed using Picard v1.130 (http://broadinstitute.github.io/picard/). Tracks were created using HOMER software v4.7 (*3*).

### WGBS analysis

Whole-genome bisulfite sequencing data were analyzed as described in Walter et al., 2016. Briefly, the first eight base pairs of the reads were trimmed using FASTX-Toolkit v0.0.13: http://hannonlab.cshl.edu/fastx_toolkit/index.html Adapter sequences were removed with Cutadapt v1.3 (*4*) and reads shorter than 16 bp were discarded. Cleaned sequences were aligned onto the mouse reference genome (mm10) using Bismark v0.12.5 (*5*) with Bowtie2-2.1.0 (*6*) and default parameters. Only reads mapping uniquely on the genome were conserved. Methylation calls were extracted after duplicate removal. Only CG dinucleotides covered by a minimum of 5 reads were conserved.

### RNA-seq analysis

2×100bp paired-end reads were mapped onto the mouse reference genome (mm10) using STAR v2.5.0a (*7*) reporting unique alignments and allowing at most 6 mismatches per fragment. Tracks were created using HOMER software v4.7 (*3*).

### 4C-seq analysis

Adapters were first trimmed using Trim Galore: v0.4.0, https://www.bioinformatics.babraham.ac.uk/projects/trim_galore/. Samples were demultiplexed using the script provided with the FourCSeq R package (v1.12.0) (*8*). Inverse primer sequences were removed on the 3′ end of the reads using Cutadapt v1.12 (*4*). Reads shorter than 15bp were discarded. Cleaned sequences were aligned onto the mouse reference genome (mm10) using Bowtie2 v2.1.0 (*6*) allowing one mismatch in the seed (22bp) and an end-to-end alignment. Subsequent steps were performed using the FourCSeq R package (v1.12.0). The mouse reference genome was *in-silico* digested using the two restriction enzymes. Restriction fragments that did not contain a cutting site of the second restriction enzyme or are smaller than 20bp were filtered out. Fragments 2.5 kb up- and downstream from the viewpoint were excluded during the procedure. Intrachromosomal contacts were kept. Valid fragments were quantified. The fragment counts were then normalized per one million reads. Data were smoothed using a running mean function with 5 informative fragments.

### Data Resources

Raw and processed sequencing data reported in this paper have been submitted to GEO, accession number pending.

---

**Figure S1.**
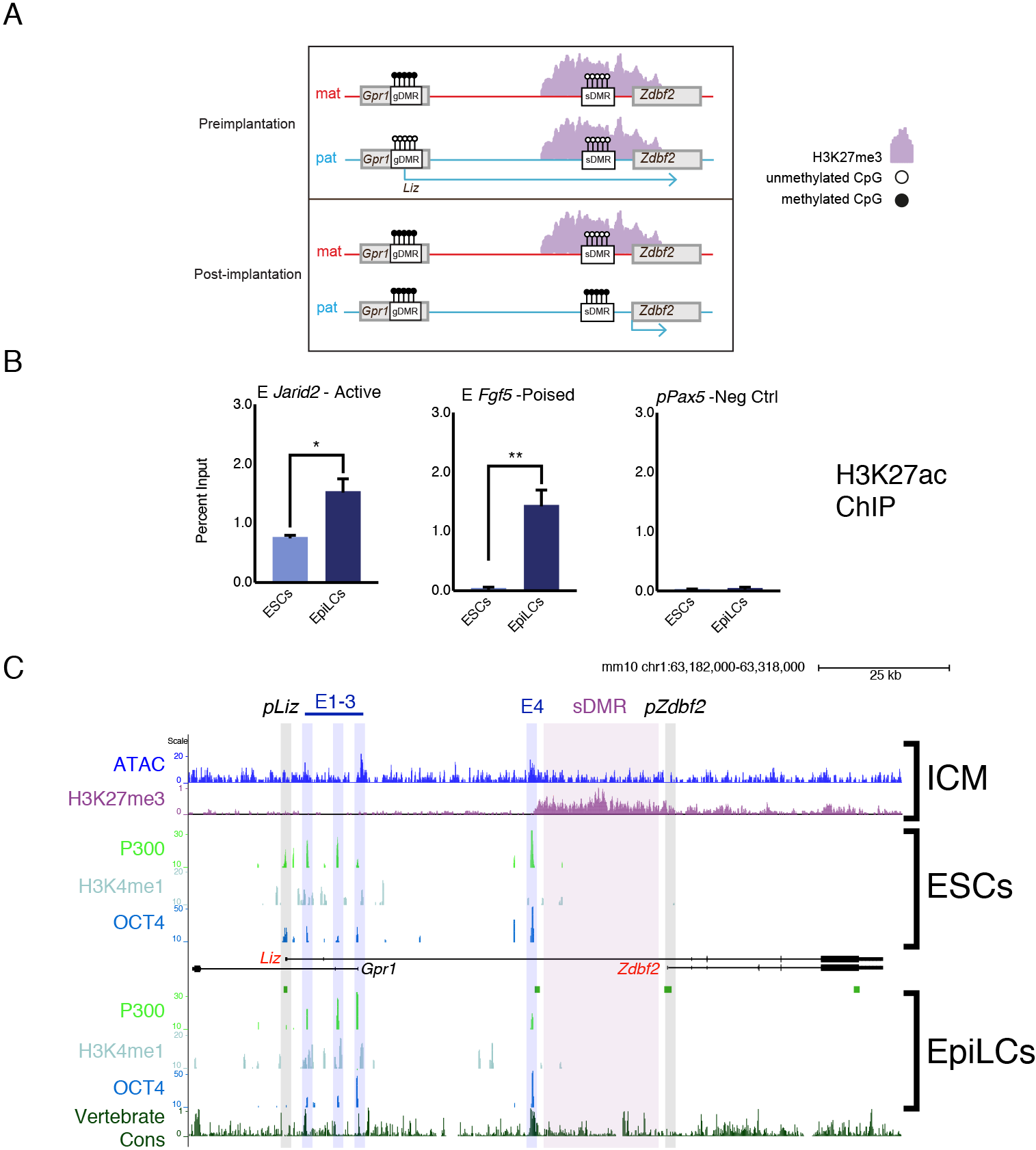
Chromatin landscape of the *Liz/Zdbf2* locus *in cellula* and *in vivo*, Related to Figure 1. **(A)** Imprinted regulation of *Liz* and *Zdbf2 in vivo.* The *Liz* promoter is DNA methylated and silent on the maternal allele. Conversely, the paternal allele is expressed, leading to *de novo* DNA methylation at the paternal sDMR, and *Zdbf2* activation. The epigenetic setting is programmed around the time of implantation in embryogenesis, but is then stably maintained throughout life. **(B)** H3K27ac ChIP-qPCR at control loci. E *Jarid2* is active in ESCs and EpiLCs, whereas E *Fgf5* is a poised enhancer (Buecker et al., 2014). *pPax5* is a negative control. Data is shown as ± s.e.m. from three biological replicates. **(C)** *In vivo* and ***in cellula*** chromatin landscape of *Liz/Zdbf2* locus. Top panel: in the *in vivo* ICM, the chromatin accessibility (Wu et al., 2016) and H3K27me3 (Liu et al., 2016) patterns resemble the *in cellula* system. Bottom panels: published data (Buecker et al., 2014) showing that active and poised enhancers are bound by P300 and OCT4. E1-3 are marked by H3K4me1, but not E4. E4 exhibits higher vertebrate conservation (PhastCons) than other enhancer elements. Statistical analyses were performed by two-tailed unpaired t-test: * P ≤ 0.05, ** P ≤ 0.01

**Figure S2.**
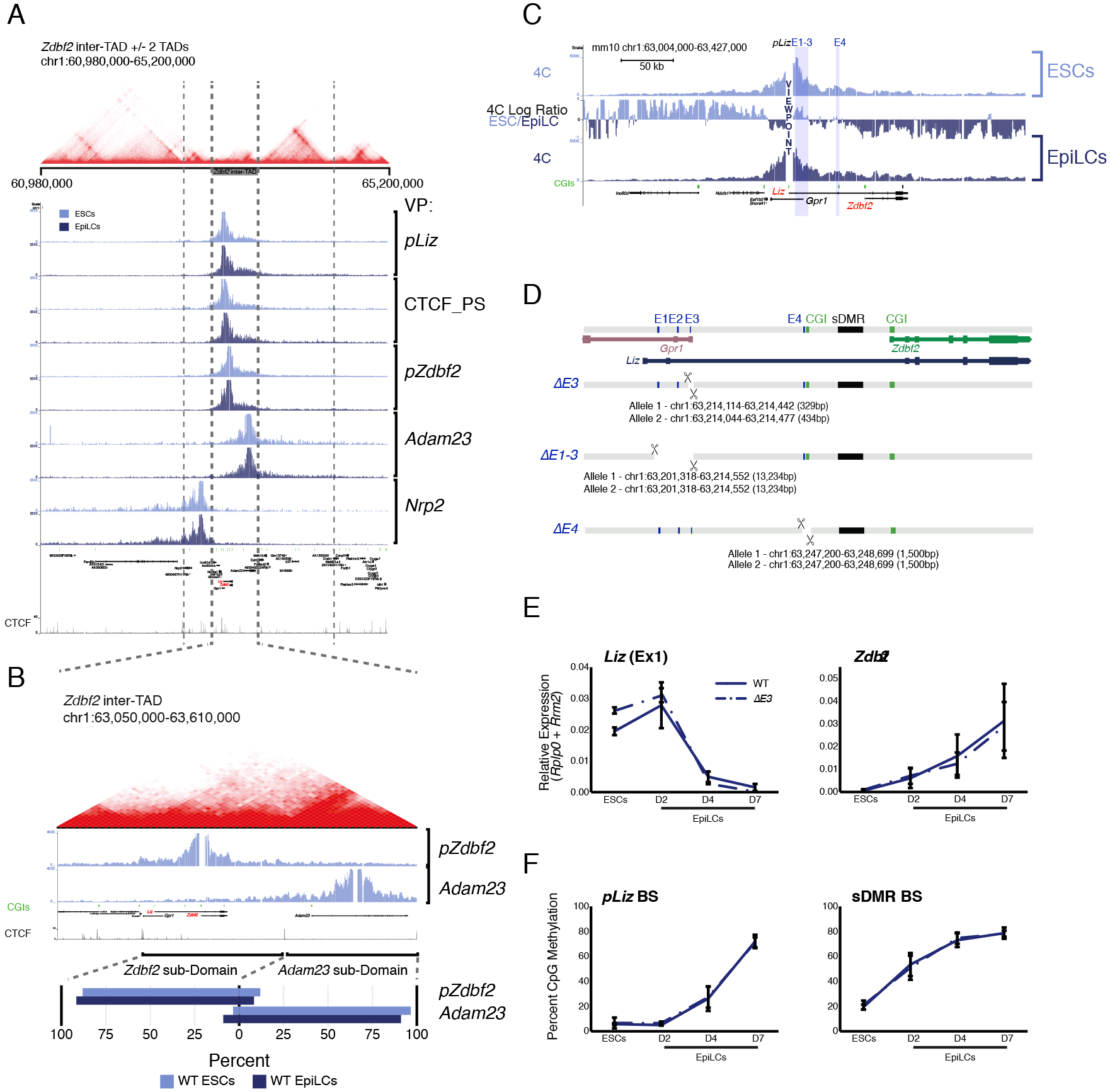
Chromosome conformation analyses of *Liz/Zdbf2* genomic region and enhancer deletions, Related to Figure 1 and Figure 2. **(A)** Hi-C data from mouse ESCs (Bonev et al., 2017) overlaid 4C-seq data from ESCs and EpiLCs (this study). *Liz/Zdbf2* is located in an inter-TAD, and 4C-seq shows that interactions in the locus are restricted to the inter-TAD. **(B)** Top: Zoom-in of inter-TAD indicates that the region can be further subdivided, with *pZdbf2* primarily interacting in a compartment within inter-TAD. Bottom: Summaries of the interaction frequencies (unnormalized reads) of *pZdbf2* and *Adam23* VPs in ESCs and EpiLCs. In both cell types, the interactions mainly occur in respective subdomains of the inter-TAD. The boundary was defined as a CTCF binding site. **(C)** 4C-seq tracks from the *pLiz* VP in ESCs and EpiLCs. Ratios between 4C-seq signals is indicated in between the samples, and gene tracks are below. In ESCs, *pLiz* exhibits increased interactions at E1-3, but interactions remain high in EpiLCs, likely due to proximity. *pLiz* also seems to be less restricted in EpiLCs, with increased interactions on “right” side of locus. The screen shot represents data from one biological replicate (two total). **(D)** Alleles generated by CRISPR/Cas9 mediated deletions of enhancer elements. **(E)** RT-qPCR of *Liz* (left) and *Zdbf2* (right) during EpiLC differentiation in WT and the Δ*E3* mutant. There is no effect on *Liz* or *Zdbf2* expression. Data are shown as ± s.e.m. from at least four biological replicates for each genotype. **(F)** DNA methylation of *pLiz* (left) and the sDMR (right) during EpiLC differentiation as measured by BS-pyro in WT and the Δ*E3* mutant. DNA methylation is unperturbed. Data are shown as ± s.e.m. from four biological replicates for each genotype.

**Figure S3.**
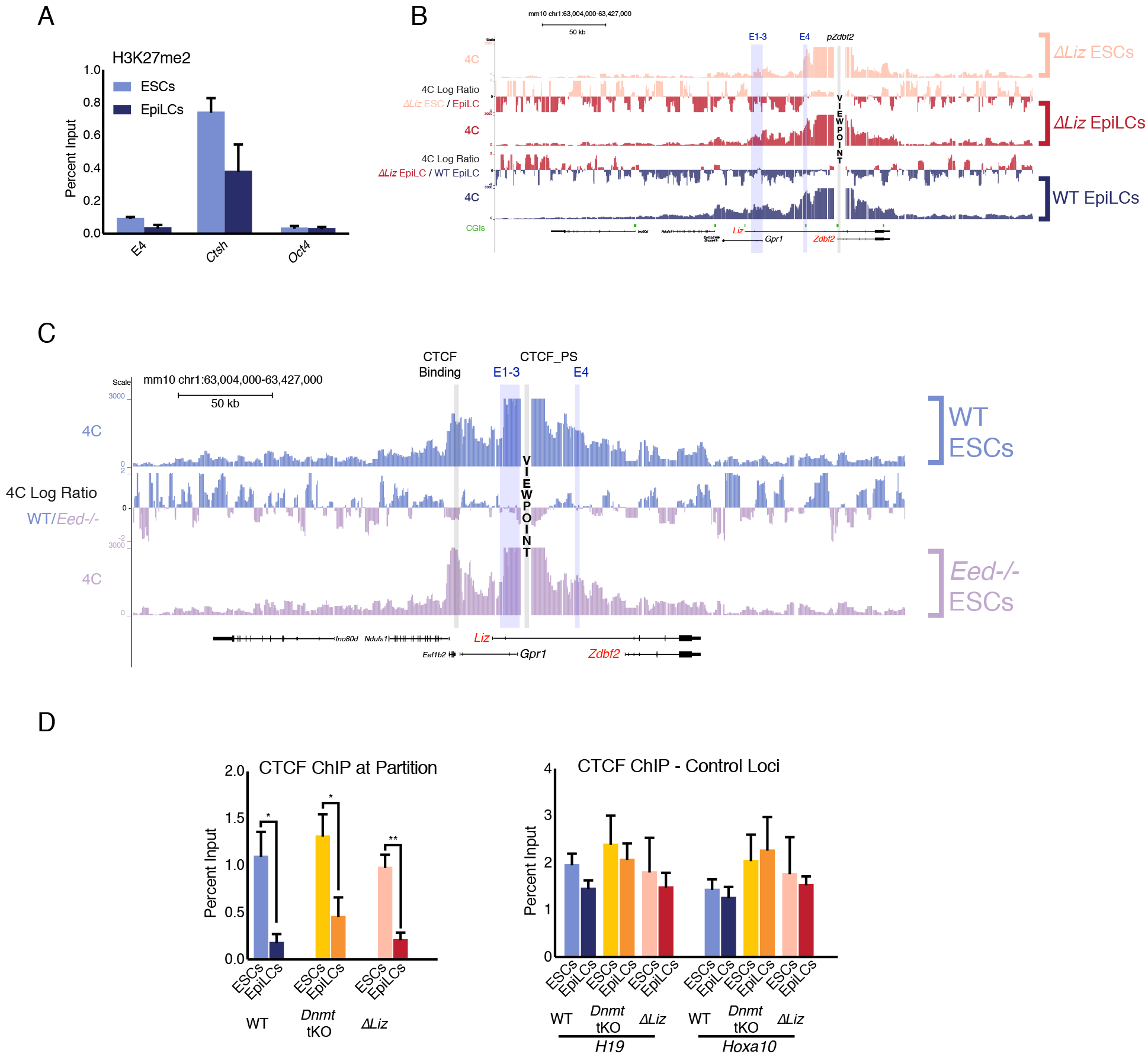
Minimal effect on chromosome conformation in *Liz* and polycomb mutants, Related to Figure 3 and Figure 4. **(A)** H3K27me2 ChIP-qPCR at E4 in WT ESCs and EpiLCs. E4 is depleted for the mark in both cell types. Data is shown as ± s.e.m. from three biological replicates. *Ctsh* and *Oct4* are positive and negative controls, respectively. **(B)** 4C-seq tracks from the *pZdbf2* VP in Δ*Liz* ESCs and EpiLCs and WT EpiLCs. Ratios between 4C-seq signals is indicated in between the samples, and gene tracks are below. While *pZdbf2* interacts with E1-4 at greater frequencies in Δ*Liz* EpiLCs compared to Δ *Liz* ESCs, it is not to the same extent as WT EpiLCs. The screen shot represents data from one biological replicate (two total). **(C)** 4C-seq tracks from the CTCF_PS VP in WT and *Eed*−/− ESCs. Ratios between 4C-seq signals is indicated in between the samples, and gene tracks are below. The partition loop still persists in *Eed*−/− ESCs, even though *Zdbf2* is active. The screen shot represents data from one biological replicate (two total). **(D)** Left panel: CTCF ChIP-qPCR in WT, *Dnmt* tKO and Δ*Liz* ESCs and EpiLCs. CTCF binding is depleted at the CTCF_PS in EpiLCs in all three contexts. Data are shown as ± s.e.m. from three biological replicates. Right panel: CTCF ChIP-qPCR in same contexts at control loci. Statistical analyses were performed by two-tailed unpaired t-test: * P ≤ 0.05, ** P ≤ 0.01

**Figure S4.**
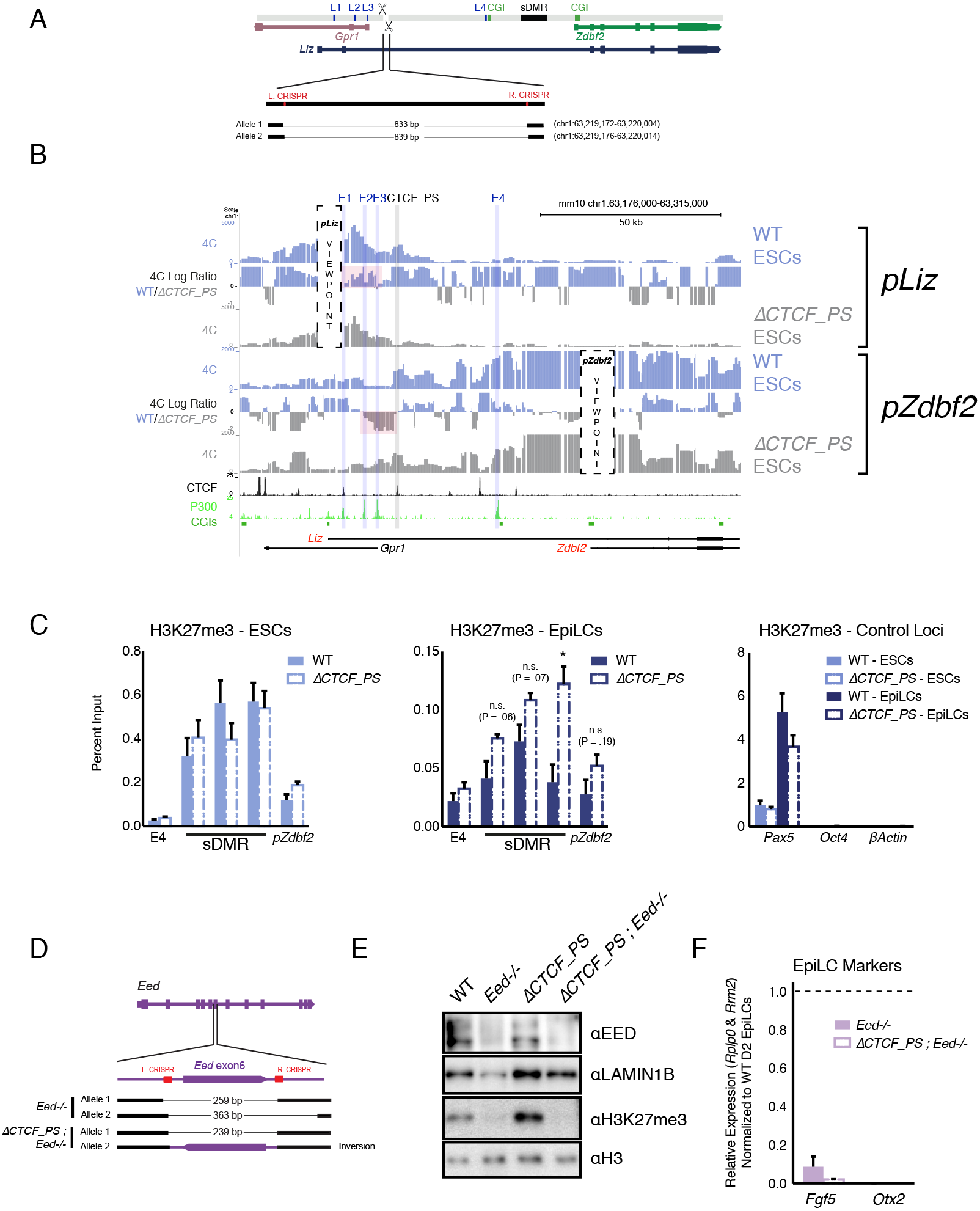
Analysis of CTCF regulation at partion site, Related to Figure 4. **(A)** Alleles generated by CRISPR/Cas9 mediated deletions of CTCF_PS **(B)** 4C-seq tracks for WT and Δ*CTCF_PS* ESCs. Top: *pLiz* VP exhibits less interactions with E1-3 in absence of CTCF partition (highlighted in pink). Bottom: *pZdbf2* VP exhibits less restricted looping with E2 and E3, although no increase with E1 (highlighted in pink) **(C)** H3K27me3 ChIP-qPCR in ESCs (Left) and EpiLCs (Middle) in WT and the Δ*CTCF_PS* mutant. There is no significant effect on polycomb dynamics in ESCs. In EpiLCs, the Δ*ACTCF_PS* mutant retains residual H3K27me3 relative to WT. H3K27me3 ChIP-qPCR in WT and Δ*CTCF_PS* ESCs and EpiLCs at control loci are displayed in right panel. Data are shown as ± s.e.m. from three biological replicates for each genotype. **(D)** Alleles generated by CRISPR/Cas9 mediated mutagenesis of *Eed* in WT and *ACTCF PS* contexts. **(E)** Western blot confirming loss of EED protein and H3K27me3 in *Eed*−/− cell lines. **(F)** RT-qPCR of in-house-generated *Eed*−/− and Δ*CTCF_PS*; *Eed−/−* lines for early epiblast markers. Values are normalized to WT EpiLCs (D2 of differentiation). The markers remain highly repressed, indicating is more likely due to polycomb deficiency, not precocious differentiation. Data are shown as ± s.e.m. from three biological replicates for each genotype. Statistical analyses were performed by two-tailed unpaired t-test: * P ≤ 0.05

**Table S1.**
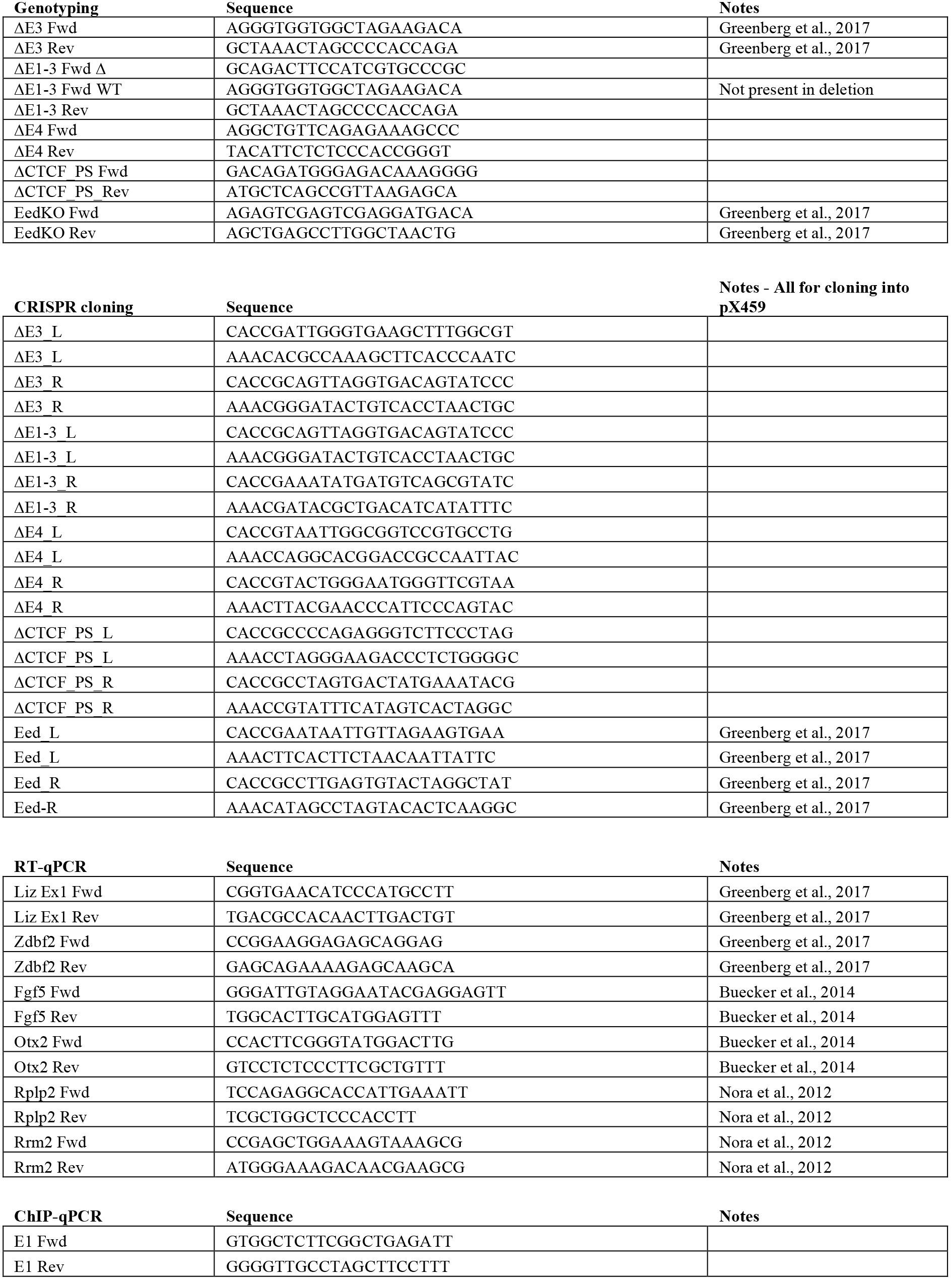
List of oligos used in this study.

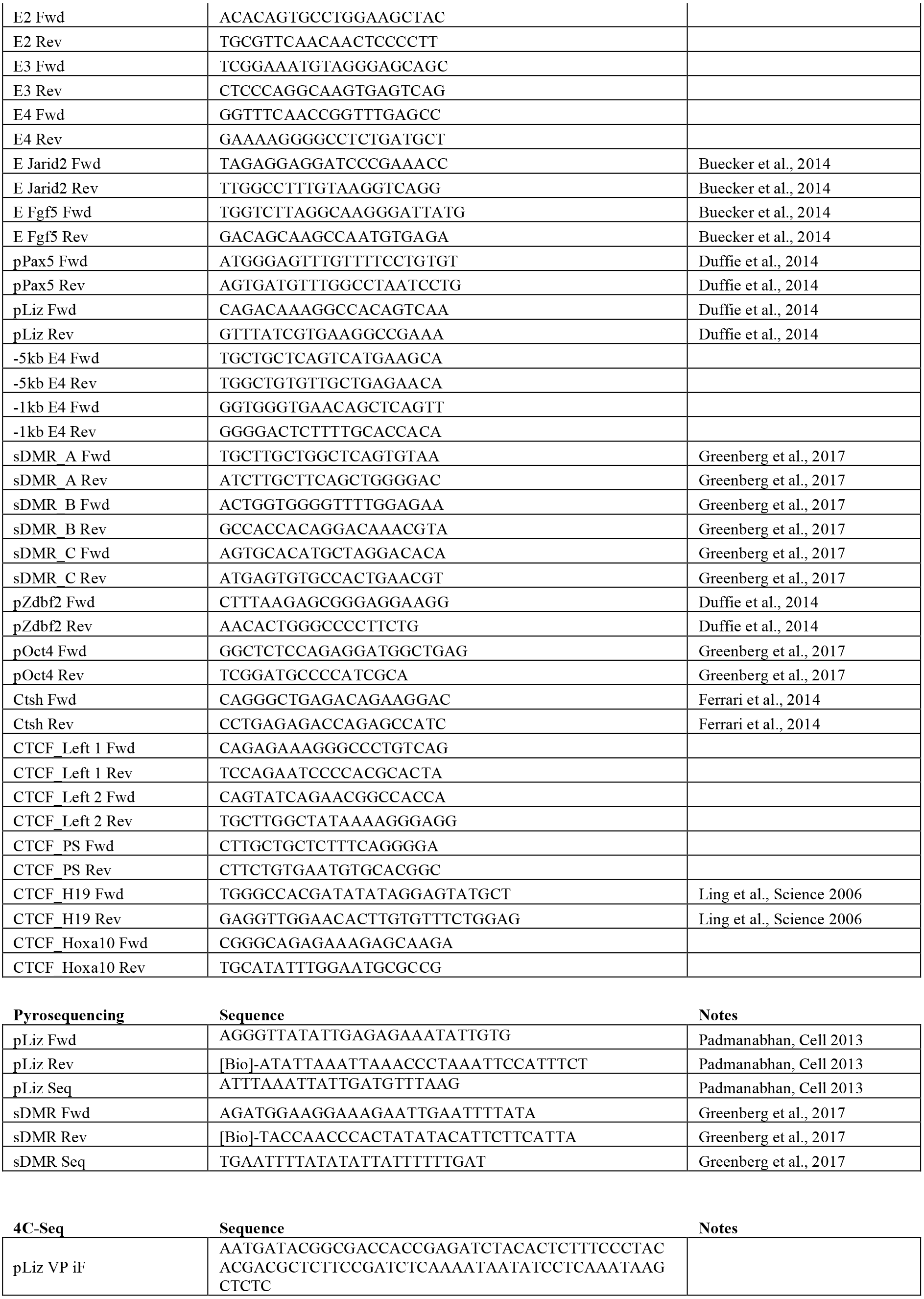

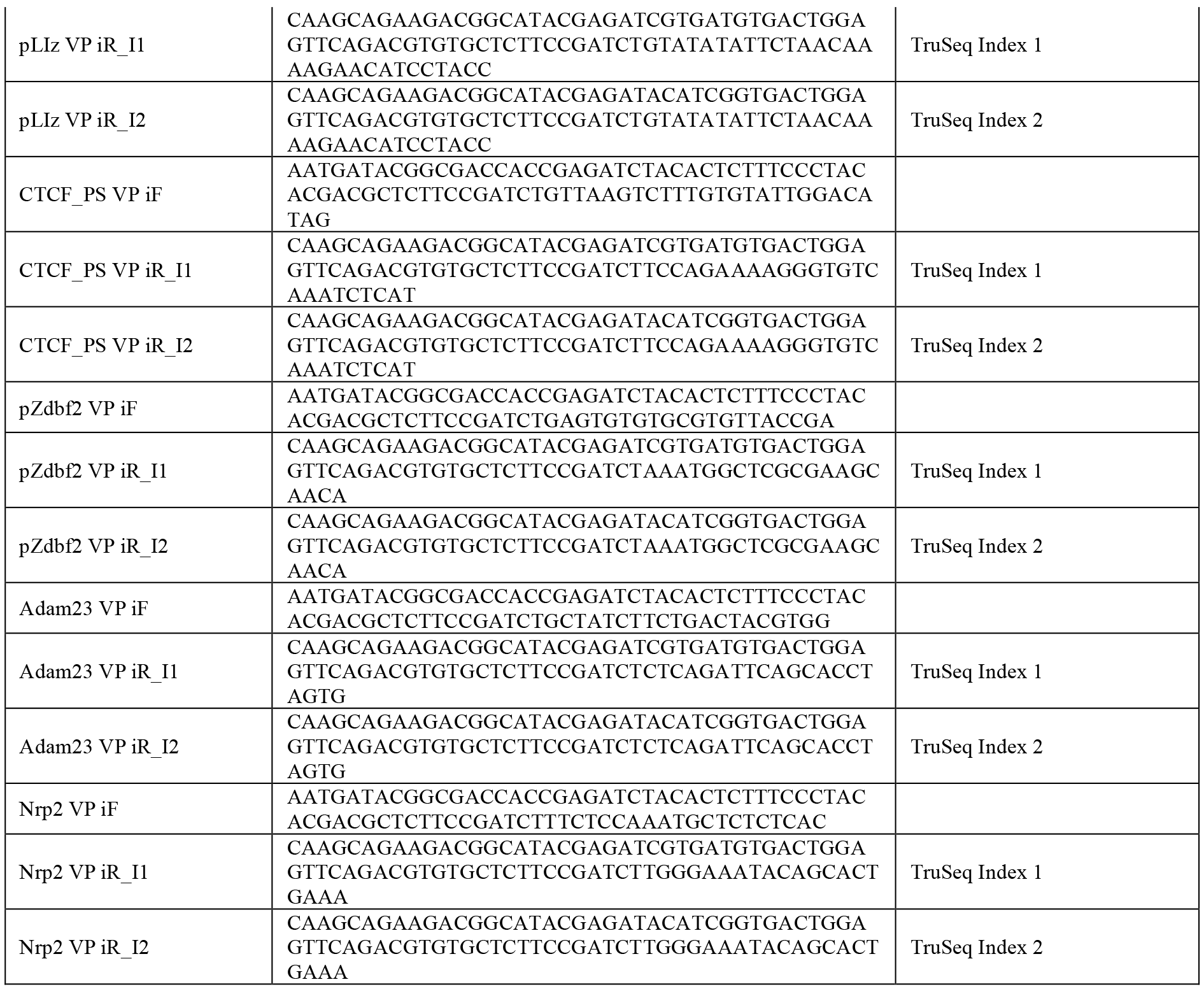

